# Proteomic and metabolic profiling reveals *APOE4*-dependent shifts in whole brain, neuronal, and astrocytic mitochondrial function and glycolysis

**DOI:** 10.1101/2025.06.15.659811

**Authors:** Colton R. Lysaker, Chelsea N. Johnson, Vivien Csikos, Edziu Franczak, Maggie Benson, Caleb A. Gilmore, Cole J. Birky, Xin Davis, Colin S. McCoin, Keith P. Smith, Patrycja Puchalska, Peter Crawford, Chad Slawson, John P. Thyfault, Paige C. Geiger, Jill K. Morris, Heather M. Wilkins

## Abstract

Apolipoprotein E (*APOE*) genetic variation is the strongest genetic risk factor for late onset Alzheimer’s disease (LOAD). Studies on *APOE* genotype dependent changes have largely focused on amyloid beta (Aβ) aggregation, disease pathology, and lipid metabolism. Recently, there has been increased interest in the relationship between metabolic function and *APOE* genetic variation. In this study, we examined how *APOE* genotype can alter metabolism in the brains of young male and female *APOE3* and *APOE4* targeted replacement (TR) mice. In combination with this, we also examined cell type-specific differences using induced pluripotent stem cell (iPSC) derived astrocytes and neurons. We found sex and genotype dependent changes to metabolism in the brains of young *APOE* TR mice. Specifically, *APOE4* mice show signs of metabolic stress and compensatory mechanisms in the brain. Using proteomics and stable isotope tracing metabolomics, we found that *APOE4* iAstrocytes and iNeurons exhibit signs of inflammation, mitochondrial dysfunction, altered TCA cycle and malate-aspartate shuttle activity, and a metabolic shift toward glycolysis. Taken together, this data indicates *APOE4* causes early changes to metabolism within the central nervous system. While this study establishes a relationship between *APOE* genotype and alterations in bioenergetics, additional studies are needed to investigate underlying mechanisms.

## Introduction

Alzheimer’s disease (AD) affects more than 6 million Americans, with growing evidence highlighting the impact of metabolic dysfunction in its progression (1). Genetic variation in apolipoprotein E (*APOE*) is a major risk factor for late onset AD (LOAD) where the ε*2* allele lowers risk, the ε*3* allele affords neutral risk, and the ε*4* allele increases risk (2). In addition to being a key regulator of lipid transport and metabolism, APOE also alters AD pathologies, including amyloid beta (Aβ). As such, *APOE4* is associated with increased Aβ deposition but accumulating evidence suggests that its influence on metabolic pathways may contribute to disease risk independently of Aβ (3–7).

Numerous studies have shown that patients with mild cognitive impairment (MCI) and AD exhibit deficits in brain energy metabolism years before disease symptom onset (8–11). These changes in brain metabolism can be influenced by *APOE* genetic variation (12–14). Work from our group and others show that postmortem AD brain from *APOE4* carriers display mitochondrial dysfunction and increased oxidative stress markers (15, 16). APOE4 has also been shown to alter the integrity of the blood brain barrier (BBB) which may influence metabolite exchange between the periphery and the central nervous system (CNS) including reduced brain glucose and lipid transport (17–19). Due to the difficulty of studying changes in humans, underlying mechanisms remain poorly understood and require further investigation.

Prior lipidomic, metabolomic, and magnetic resonance spectroscopy (MRS) studies in mouse models have shown that genetic variation in *APOE* can modify brain metabolite levels (20–23). For example, *APOE4* humanized mice show altered brain levels of palmitate, succinate, lactate, and malate (21). Pathways and proteins associated with oxidative phosphorylation, glycolysis, and lipid metabolism also showed altered brain expression profiles in *APOE4* humanized mice (21, 22, 24). Additionally, microglia and astrocytes which have been extensively studied *in vivo*, show *APOE* genotype dependent immune and inflammatory metabolic changes (22, 25, 26). However, most prior studies have not examined sex differences or potential early metabolic changes.

Studies performed *in vitro* have shown that *APOE* genetic variation can alter cellular metabolic profiles in distinct cell types including astrocytes, microglia, and neurons (27–30). One group found that *APOE* genetic variation can alter lipid metabolism and fatty acid coupling in primary mouse neurons and astrocytes (31). Another study found that APOE4 can drive changes to lipid trafficking in primary mouse astrocytes (32). These changes are not limited to lipid metabolism and others have found that glucose metabolism in microglia and astrocytes is altered by *APOE* genotype in both primary cell culture and induced pluripotent stem cell (iPSC)-derived models (22, 24, 33–35). Neurons have often been understudied in relation to the effects of APOE on metabolism due to their low expression of APOE under homeostatic conditions (36, 37). In these limited studies, *APOE* genetic variation has been found to alter neuronal health and metabolism (30, 31).

In this study, we employed both *in vivo* and *in vitro* models in combination with omics-based methods to interrogate *APOE* genotype-dependent changes to CNS metabolism. Here, we performed detailed metabolic profiling in both iPSC-derived neurons and astrocytes using respiration analysis, proteomics, and glucose uptake/utilization with stable isotope tracing untargeted metabolomics (ITUM). We also examined whole brain and isolated mitochondria proteomics and mitochondrial function in young *APOE3* and *APOE4* targeted replacement (TR) mice. We found that *APOE* genotype caused extensive changes to metabolism consistently across *in vivo* and *in vitro* models. This includes sex specific changes in *APOE* TR mice. These results highlight the importance of APOE in driving metabolic alterations across the brain and between specific cell types.

## Methods

### Animals

Male and female homozygous *APOE3* (B6.129P2-*Apoe^tm2(APOE*3)Mae^* N8) and *APOE4* (B6.129P2-*Apoe^tm3(APOE*4)Mae^* N8) targeted replacement (TR) mice were purchased from Taconic Biosciences (Germantown, NY) for this study (n=8 per group for genotype and sex). These *APOE* TR mice were previously generated by removing and replacing the endogenous mouse *Apoe* gene with the human *APOE* gene (38, 39). Mice were maintained on a standard chow diet (Teklad Global Rodent Diet®, 8604) for the entire study and housed at ∼25°C with a standard 12-hr light/dark cycle. Mice were aged to 4 months before the administration of 90 mg/kg ketamine and 10 mg/kg xylazine followed by euthanasia. Brain tissue was collected immediately following euthanasia and used for mitochondrial isolation or stored at -80°C. The mice used in this study were also used in another publication (40). Animal work and protocols were conducted under approval from the Institutional Animal Care and Use Committee at the University of Kansas Medical Center.

### Human induced pluripotent stem cell culture

CRIPSR-Cas9 gene edited isogenic human induced pluripotent stem cells (iPSCs) homozygous for *APOE3* or *APOE4* were obtained from The Jackson Laboratory (JAX). These iPSCs were generated as previously described from the KOLF2.1J parental cell line (41). Two pairs of isogenic iPSCs were used in this study, with Pair A consisting of JIPSC001162 (*APOE3* REV/REV) and JIPSC001150 (*APOE4* SNV/SNV), and Pair B consisting of JIPSC001268 (*APOE3* REV/REV) and JIPSC001142 (*APOE4* SNV/SNV). The iPSC lines were maintained on a feeder free system using Matrigel (Corning) and mTeSR™ Plus medium (STEMCELL Technologies). Cells were passaged with ReLeSR™ (STEMCELL Technologies) when colonies became large and confluent to avoid spontaneous differentiation as outlined in manufacturer protocols.

### Generation of iNeurons and iAstrocytes

iPSCs were differentiated into neural progenitor cells (NPCs) using STEMdiff™ Neural Induction Medium (NIM) from STEMCELL Technologies. iPSCs were placed into a single cell suspension in NIM with SMADi/ROCKi (SMAD inhibitor and 10μM ROCK Inhibitor) in an AggreWell 800 plate. Embryoid bodies were cultured in the AggreWell™ plate for 5 days with NIM/SMADi partial medium changes daily. Embryoid bodies were plated onto Matrigel (Corning) coated plates and fed daily with NIM/SMADi medium until day 12 to allow neural rosette formation. Neural rosettes were selected using Neural Rosette Selection Reagent (STEMCELL Technologies) and plated onto Matrigel (Corning) coated dishes with NIM/SMADi. The medium was changed daily for 7 days, after which NPCs were cryopreserved and split into defined STEMdiff™ Neural Progenitor Medium (NPM) (STEMCELL Technologies). For forebrain neuronal differentiation, NPCs were plated on PLO/Laminin (Sigma) coated dishes and cultured in NPM. The following day the medium was changed to STEMdiff™ Forebrain Neural Differentiation Medium (STEMCELL Technologies), which was changed daily for 5 days. Cells were then plated onto PLO/Laminin coated dishes in defined BrainPhys™ Medium (with 1X N2A supplement, 1X SM1 supplement, 20 ng/mL BDNF, 20 ng/mL GDNF, 1mM cAMP, and 200 nM ascorbic acid) for neuronal maturation. Neurons were matured for a minimum of 14 days before use in downstream experiments. For astrocyte differentiation, NPCs were plated onto Matrigel coated dishes in NPM. The following day cells were placed in astrocyte differentiation medium consisting of DMEM containing 4.5 g/L D-glucose, 1X B27, 1% FBS, 4mM glutamine, 8 ng/mL bFGF, 5ng/mL CNTF, 10 ng/mL BMP8, 10 ng/mL Activin A, 10 ng/mL heregulin 1b, and 200 ng/mL IGF1. Medium was changed every other day and cells were passaged as needed. After 30 days or approximately 5-6 passages astrocytes were used in experiments.

### Mitochondrial isolation from whole brain

Upon collection, left brain hemispheres were immediately placed in 8 mL of ice-cold mitochondrial isolation buffer (MIB) (225 mM mannitol, 75 mM sucrose, 6 mM K2HPO4, 1 mM EDTA, 0.1% fatty acid free BSA, pH 7.2) and homogenized using a Teflon pestle on ice. Homogenates were then centrifuged at 1,500 x g for 10 min at 4°C to remove cell debris. The supernatant was collected and centrifuged at 8000 x g for 10 min at 4°C. The pellet was then resuspended on ice in 6 mL of MIB and centrifuged again at 6,000 x g for 10 min at 4°C. The pellet was resuspended in 500 μL of ice-cold MiR05 buffer (0.5 mM EGTA, 3mM MgCl_2_, 60 mM KMES, 20 mM glucose, 10 mM KH_2_PO_4_, 20 mM HEPES, 110 mM sucrose, 0.1% BSA, pH 7.1). A BCA assay (ThermoFisher) was performed to determine protein content. Mitochondrial samples were then either immediately used for respiration analysis or an aliquot was immediately frozen at -80°C for proteomics.

### Proteomics

Whole brain samples for proteomics were prepared by crushing flash frozen tissue on liquid nitrogen with a pestle. Roughly 80-100 mg of brain tissue was then placed into new tubes and resuspended in homogenization buffer (1% Triton X-100, 50 mM HEPES, 12 mM Na pyrophosphate, 100 mM NaF, 10 mM EDTA, protease inhibitor, and phosphatase inhibitor). Samples were then lysed in a TissueLyser II (QIAGEN) with a stainless-steel metal bead. Freshly lysed whole brain lysates were centrifuged at 15,000 x g for 25 min at 4°C. To determine protein content a BCA assay was used before diluting samples in fresh lysis buffer to a concentration of 1 ug/uL (100 ug total). Protein content was determined from previously frozen isolated liver mitochondria samples which were diluted to 1 μg/μL (100 μg total).

Neuron and astrocyte cell lysates were prepared by first placing culture plates on ice and washing with ice cold 1X PBS. Cells were then scraped with 1X PBS containing protease inhibitors (ThermoFisher, A32965) and centrifuged at maximum speed for 5 min. The supernatant was removed, and the pellet was resuspended and lysed in 200 uL RIPA (Millipore) with protease inhibitors (ThermoFisher, A32965). After a BCA assay was performed, cell lysates were diluted to 0.25 ug/uL (50 ug total of protein).

Samples were sent to the University of Arkansas Medical Center (UAMS) for proteomics. Briefly, protein samples were extracted using chloroform/methanol phase separation with a subsequent trypsin digestion to acquire peptides. Whole brain and isolated mitochondria were analyzed using liquid chromatography mass spectrometry (LC/MS) on a Orbitrap Exploris™ 480 (ThermoFisher) with data independent acquisition (DIA). Astrocyte and neuron samples were analyzed using a Orbitrap Astral (ThermoFisher) with DIA.

### Stable isotope tracing untargeted metabolomics

Cells were grown in 6-well dishes coated with PLO/Laminin (neurons) or Matrigel (astrocytes); 2 wells of a 6 well plate constituted one sample. Prior to labeling, cells were starved in glucose, phenol, and serum free DMEM for 1 hour. After removal of the media, cells were treated with labeled [U-^13^C_6_]-glucose (Cambridge Isotope) or [^12^C]-glucose (Sigma) in serum and phenol free DMEM for 24 hours at a concentration of 2.5 mM for neurons and 25 mM for astrocytes. All cell incubations with media took place at 37°C, 5% CO_2_. Media was collected and flash frozen in liquid nitrogen. Cells were washed 1X with warmed PBS and 1X with warmed water. Cell plates were then submerged in liquid nitrogen for 30 seconds. 500 µL ice cold methanol was added to each well and cells were scraped, transferred to a microfuge tube and stored at -80°C. The supernatant was removed using a SpeedVac prior to the analysis.

Metabolite extraction and LC/MS analysis was performed as described previously (42–44). An extraction was performed on the cells using 1 mL of -20°C ice cold methanol:acetonitrile:H_2_O at a ratio of 2:2:1 (v/v/v). Three subsequent cycles of vortexing, freezing-thawing followed by water bath sonication were performed. Following this, samples were incubated for 1 hour at -20°C with a subsequent centrifugation for 10 min at maximum speed. The supernatant was transferred to new tubes and supernatant evaporated with SpeedVac. The final pellet was reconstituted with 40 μL of acetonitrile(ACN):H_2_O at a ratio of 1:1 (v/v) before being vortexed and incubated for 1 hour at 4°C. LC/MS analysis was then performed after centrifuging the supernatant.

We performed LC analysis with a Dionex Ultimate 3000 RSLC column (100 mm x 1 mm, 3 um particle size). The column was used in hydrophilic interaction liquid chromatography (HILIC) mode. Two mobile phase compositions were used which included A (95% H_2_O, 5% ACN, 10 mM NH_4_OAc/NH_4_OH, pH 9.5) and B (95% ACN, 5% H_2_O, 10 mM NH_4_OAc/NH_4_OH, pH 9.5. Extracts were run and separated on a Luna NH_2_ with a binary gradient 75-0% B for 45 min, then 0% B for 12 min, and 75%B for 13 min at 50 μL/min. The column temperature was kept a 30°C with a 4 uL injection volume. A Thermo Q Exactive Plus with heated ESI source was used to perform MS. The MS was run in negative mode with the resolution set at 70,000, the AGC target to 3e6 ions, and a 200 ms maximum injection time. The mass scan for extract analysis was 68-1020 m/z. The following ESI parameters were used: auxiliary gas 10, sweep gas 1, spray voltage -3 kV, capillary temperature 275°C, S-lens RF 50, and auxiliary gas temperature 150°C. The sheath gas flow was set to 35 (AU) for extract analysis.

Analysis of metabolites was completed using a group of standard compounds and matching identified metabolites with their retention times. Raw data was converted to a mzXML format using MSConvert with the vendor peak-picking option selected. These files were then analyzed using X^13^CMS R package as previously described (45, 46). Data was processed with R studio XCMS (v. 1.4) package to pick peaks (method = ‘centWave’, ppm = 2.5, peak width = c(20, 180) and align retention times (bw=10, mzwid=0.015, retention time correction method = ‘obiwarp’) across samples. Parameters used for X^13^CMS included RTwindow = 10, ppm = 5, noiseCutoff = 10000, intChoice =“intb”, alpha = 0.05 within getIsoLabelReport().

### Western blotting

Whole cell lysates from astrocytes and neurons were collected on ice and lysed using RIPA buffer (Millipore) with protease inhibitors (ThermoFisher, A32965). A BCA assay (ThermoFisher) was performed to normalize protein loading, and 5 ug/well of protein was resolved on 4-15% Criterion TGX gels (Bio-Rad). Gels were transferred to PVDF membranes (Bio-Rad) and blocked for 1 hour at room temperature with 5% bovine serum albumin (BSA) in 1X phosphate buffered saline with Tween 20 (PBST). Blots were incubated overnight at 4°C with primary antibody diluted in 1X PBST with 5% BSA. After three washes in 1X PBST, blots were incubated at room temperature for 1 hour with secondary antibody diluted in 5% BSA in 1X PBST. Imaging was performed using West Dura ECL substrate (ThermoFisher) on a ChemiDoc XRS imaging system (Bio-Rad) with automatic exposure. O-GlcNAc primary antibody from ThermoFisher (MA1-072) was diluted 1:000. A goat anti-mouse IgG HRP conjugated secondary antibody (Bio-Rad, #1706516) was used at a dilution of 1:5000.

### Immunocytochemistry

Cells were fixed using 4% paraformaldehyde (PFA) for 15 min. Cells were washed three times with 1X PBS (Gibco) before being permeabilized with 0.1% Triton X-100 (Sigma) for 10 min in 1X PBST. Cells were then washed three times with 1X PBS (Gibco) and blocked for 1 hr at room temperature with 1% BSA in 1X PBST. The blocking solution was removed, and cells were incubated overnight with primary antibodies at 4°C. The following day cells were washed with 1X PBS (Gibco) three times followed by a 1 hr incubation at room temperature with secondary fluorophore conjugated antibodies. The nuclear counterstain Hoechst was used at a concentration of 2 μg/mL for 30 min. Finally, cells were washed three times with 1X PBS (Gibco) before being imaged on a BioTek Cytation 1 (Agilent). Primary antibodies included S100B diluted 1:100 (Abcam, ab52642) and MAP2 diluted 1:200 (Abcam, ab32454). We used DyLight™ conjugated secondary antibodies (ThermoFisher) at a concentration of 2 μg/mL.

### Isolated mitochondria respiration

Brain mitochondrial oxygen consumption rates (OCR) were measured using an Oroboros O2k respirometer. Isolated mitochondria were added to the chamber and state 2 respiration rate was determined using 5 mM pyruvate and 5 mM malate. We determined state 3 respiration with the addition of with 4 mM ADP. State 3G and state 3S were measured after the addition of 2mM glutamate and 10 mM succinate, respectively. The sequential addition assessed maximal respiratory flux with carbonyl cyanide-p-trifluoromethoxy phenylhydrazone (FCCP, Uncoupled, 4 µM). Respiration values were normalized to total protein amount loaded and across days of data acquisition.

### Cellular respiration

A Seahorse XF Pro Analyzer (Agilent) was used to determine cell OCR and extracellular acidification rates (ECAR). Astrocytes were plated at a density of 30,000 per well on Matrigel (Corning) coated Seahorse cell culture plates. Neurons were cultured on PLO/Laminin coated plates at a density of 40,000 cells/well. Substrates were successively injected to measure OCR or ECAR for different respiration rates. *Mitochondrial Stress Test:* seahorse base medium contained 25 mM glucose (astrocytes) or 2.5 mM glucose (neurons), 2 mM glutamine, with injections A) 2 µM oligomycin B) and C) 0.5 µM FCCP D) 1 µM antimycin A and rotenone. *Glycolysis Stress Test*: Seahorse base medium contained 2 mM glutamine with injections A) 10 mM glucose B) 2 µM oligomycin C) 100 mM 2-deoxy-glucose (2-DG). *Electron Transport Chain Flux*: Medium was 1X MAS (220 mM mannitol, 70 mM sucrose, 10 mM KH_2_PO_4_, 2 mM HEPES, 5 mM MgCl_2_, 1 mM EGTA, 0.2% fatty acid free BSA) supplemented with 10 mM pyruvate/5 mM malate/4 mM ADP and 1 nM plasma membrane permeabilization (PMP) reagent with injections A) 10 mM succinate/1 µM rotenone B) 1 µM antimycin A C) 0.5 mM TMPD/1 mM ascorbate D) 50 mM azide. *Fuel Dependency/Capacity:* To determine glutamine dependency, baseline OCR measurements were taken followed by injections of A) 3 µM BPTES and B) 4 µM Etomoxir with 2 µM UK5099. To determine glutamine capacity baseline OCR measurements were taken followed by injections of A) 4 µM Etomoxir with 2 µM UK5099 and B) 3 µM BPTES. To determine fatty acid dependency baseline OCR measurements were taken followed by injections of A) 4 µM Etomoxir and B) 3 µM BPTES with 2 µM UK5099. To determine fatty acid capacity baseline OCR measurements were taken followed by injections of A) 3 µM BPTES with 2 µM UK5099 and B) 4 µM Etomoxir. To determine glucose dependency baseline OCR measurements were taken followed by injections of A) 2 µM UK5099 and B) 4 µM Etomoxir with 3 µM BPTES. To determine glucose capacity baseline OCR measurements were taken followed by injections of A) 4 µM Etomoxir with 3 µM BPTES and B) 2 µM UK5099.

OCR and ECAR were measured following a 3-min mix after the addition of injections and measured over 3 min three times. All data were analyzed using the Agilent Seahorse Wave software and Microsoft Excel.

### Mitochondrial indicator assays

Astrocytes and neurons were plated at a density of 2.0-3.0x10^4^ cells/well of a transparent bottom 96 well plate (3603, Corning). To assess mitochondrial membrane potential we used 200 nM tetramethyl rhodamine, ethyl ester (TMRE) (ThermoFisher). Mitochondrial superoxide was measured with 500 nM MitoSOX™ Red (ThermoFisher) and hydrogen peroxide (H2O2) was determined using 50 μM Amplex™ Red (ThermoFisher) with 0.1 U/mL horseradish peroxidase (HRP). Levels of mitochondrial calcium (Ca2+) were measured using 1 μM Rhod-2, AM (ThermoFisher). Hoechst (ThermoFisher) was used for normalization at a concentration of 10 μg/mL. Cells were stained with respective dyes diluted for 30 min at 37°C and then washed twice with 1X Hanks’ Balanced Salt Solution (HBSS) (Gibco). Fluorescence intensity for each dye was read with a Tecan Infinite 200 PRO® and normalized to Hoechst.

### Proteomic data analysis

Data analysis of samples was completed using Spectronaut® software (Biognosys version 18.3). UniProt databases were used to identify both mouse (Mus musculus) and human (Homo sapiens) proteins. Quality control (QC) and normalization was carried out with proteiNorm (47). MitoCarta 3.0 was used to further identify mitochondrial specific proteins in the isolated brain mitochondria samples (48). Proteins with a p-value < 0.05 were considered significant. QIAGEN Ingenuity Pathway Analysis (IPA) was used for pathway and comparison analysis. Proteins included in pathway analysis had a p-value cutoff of < 0.2 for whole brain and isolated mitochondria samples. For astrocytes and neurons, pathway analysis was performed on proteins with a p-value < 0.05 and a logFC < -0.5 and > +0.5. Pathways with a z-score of ≤ -2 and ≥ +2 and a log fold change (logFC) of ≥ 1.3 were considered significant. Proteins in specific cellular functions and pathways were identified by cross referencing data with the Kyoto Encyclopedia of Genes and Genomes (KEGG) and IPA.

### Statistical analysis

Values are shown as mean ± SD. Data analysis was completed using GraphPad Prism 10 software and Microsoft Excel. Outliers were identified and removed from datasets using Grubbs’ test with a significant alpha of 0.05. Significance for simple comparisons was calculated using an unpaired t test. For comparisons of multiple groups we used a two-way analysis of variance (ANOVA) followed by Fisher’s Least Significant Difference (LSD) post-hoc test if significant interactions were identified. A cutoff p-value < 0.05 was considered significant for all analyses. Pearson correlation analysis was used to assess correlations. All graphs and figures were generated using GraphPad Prism 10 software and BioRender.com.

## Results

### The whole brain proteome is altered by APOE genetic variation in young mice

Four-month-old male and female *APOE3* or *APOE4* targeted replacement (TR) mice were used to examine whole brain and isolated mitochondrial proteomics (n=3 per genotype/sex) in addition to Oroboros O2k respiration analysis (n=8 per genotype/sex) (**Figure 1A**). We first examined whole brain protein expression of APOE and observed reduced levels in *APOE4* mice (**Figure 1B**). Whole brain proteomics revealed 348 differently expressed (DE) proteins between female *APOE4* and *APOE3* mice versus 326 between male *APOE4* and *APOE3* mice (**Figure 1C**). Compared to female *APOE3* mice *APOE4* females had increased expression of proteins involved in mitochondrial function and bioenergetics (mt-Co3, Sdhd, Adcy3, Slc2a1/Glut1), ubiquitination/inflammation (Otulin), peptidase activity (Serpina1e), and clathrin recruitment/receptor sorting (Ap1s2). There was reduced expression of proteins involved in ketone metabolism (Hmgcs2), protein quality control (Selenof), Jak/Stat and tyrosine receptor signaling (Jak1), glutamate receptor signaling (Grik5), coenzyme Q synthesis (Coq3), and centriole duplication (Mdm1) (**Figure 1D**). IPA revealed a significant upregulation of protein sorting signaling, neutrophil degranulation, neddylation, transport of ions, and RAB geranylgeranylation pathways in *APOE4* female mice when compared to their *APOE3* counterparts (**Figure 1E**). Down regulated pathways in *APOE4* females included SRP-dependent co-translational protein targeting, response to amino acid deficiency, non-sense mediated decay, processing of mRNA, and mitochondrial translation pathways (**Figure 1E**).

**Figure 1.**
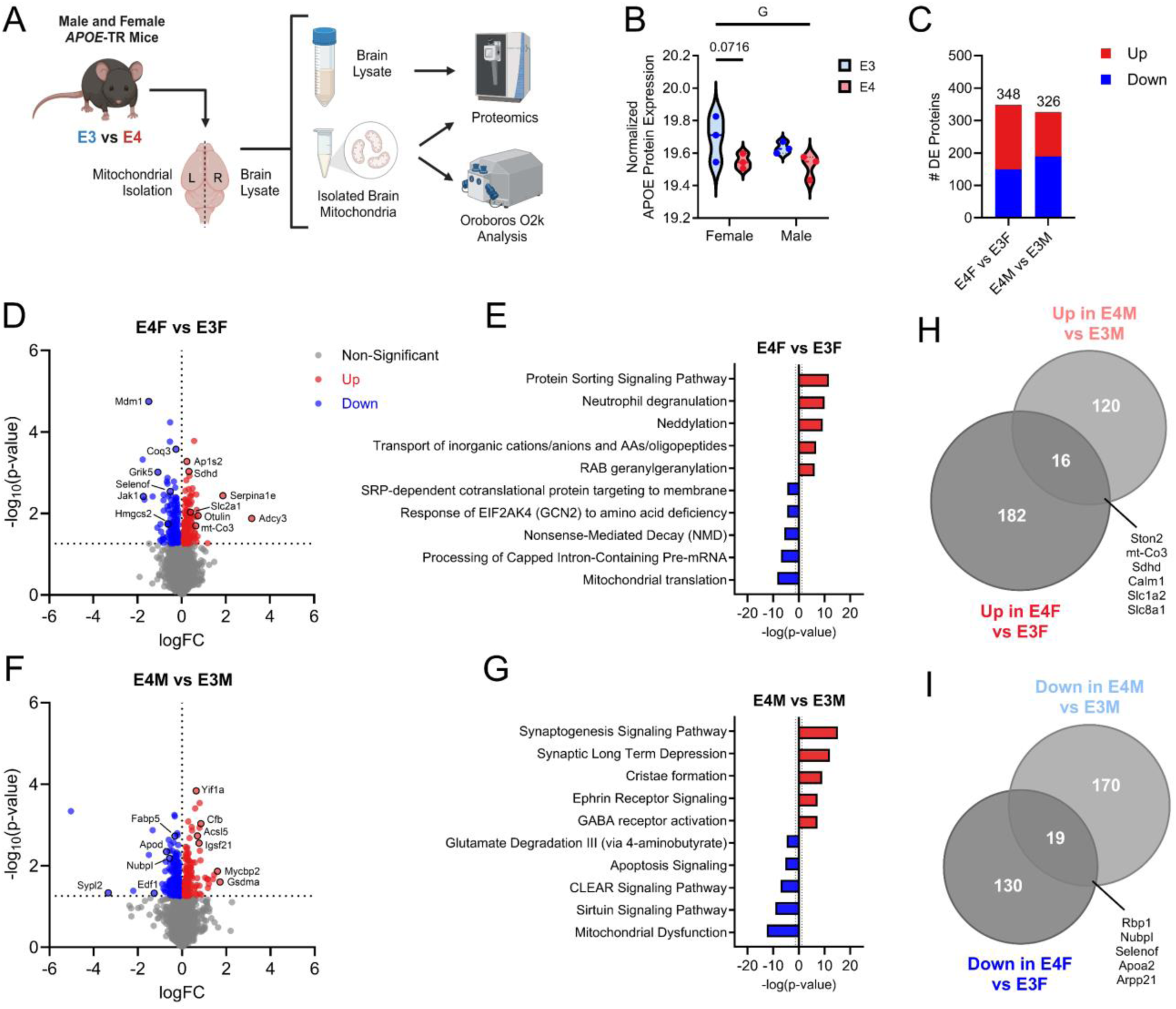
Whole brain proteomics in male and female *APOE3* and *APOE4* TR mice. A. Study schematic showing male and female *APOE3* and *APOE4* targeted replacement (TR) mice. Whole brain lysate or isolated brain mitochondria were used for proteomic (n=3 per group). B. Violin plot of APOE protein expression from whole brain in male and female *APOE3* and *APOE4* mice. C. The number of differentially expressed (DE) proteins in male and female *APOE4* vs. *APOE3* mice. D. Volcano plot showing up and down regulated proteins in female *APOE4* vs. *APOE3* mice. E. IPA showing up and down regulated pathways in female *APOE4* mice. F. Volcano plot showing up and down regulated proteins in male *APOE4* vs *APOE3* mice. G. IPA showing up and down regulated pathways in male *APOE4* mice. H. Venn diagram of upregulated protein overlap in male and female *APOE4* vs. *APOE3* mice. I. Venn diagram of down regulated protein overlap in male and female *APOE4* vs. *APOE3* mice. G=main effect of genotype.

When compared to their *APOE3* counterparts, male *APOE4* mice had increased expression of proteins involved in fatty acid metabolism (Acsl5), endoplasmic reticulum (ER)/Golgi maintenance (Yif1a), complement and cell death/inflammation (Cfb, Gsdma), and neuronal development/inhibitory presynaptic differentiation (lgsf21, Mycbp2). There was reduced expression of proteins involved in lipid metabolism (Fabp5, Apod), endothelial cell function/VEGF signaling (Edf1), iron/sulfur assembly (Nubpl), and synaptophysin like/obesity linked proteins (Sypl2) (**Figure 1F**). Pathways significantly upregulated in *APOE4* males included synaptogenesis, synaptic long-term depression, cristae formation, ephrin receptor signaling, and GABA receptor activation (**Figure 1G**). Down regulated pathways included glutamate degradation, apoptosis, CLEAR signaling, sirtuin signaling, and mitochondrial dysfunction pathways in *APOE4* male mice when compared to *APOE3* (**Figure 1G**).

We examined the overlap of protein expression changes between male and female *APOE4* versus *APOE3* TR mice. Male and female *APOE4* mice shared 16 significantly upregulated proteins and 19 down regulated proteins when compared to *APOE3* mice (**Figure 1H and I**). Interestingly, mt-Co3, Sdhd, and Slc1a2 were upregulated in APOE4 mice (**Figure 1H**). We also observed a shared decrease of Nubpl and Apoa2 (**Figure 1I**). These findings highlight not only sex specific effects of *APOE* genotype on brain proteomics, but also shared changes in both male and female *APOE4* mice. Sex specific comparisons were further characterized in **Supplemental Figure 1**.

### APOE genotype alters respiration and the proteome of isolated brain mitochondria

To examine mitochondrial function between *APOE4* and *APOE3* mice, we leveraged Oroboros O2k respirometry analysis in isolated brain mitochondria. Carbohydrate driven respiration showed a main effect for genotype and an interaction between sex and genotype for state 3 respiration. Male *APOE3* mice had increased state 3 respiration compared to male *APOE4* mice (**Figure 2B**). No change was observed for state 2 respiration (**Figure 2A**). Carbohydrate driven respiration showed an interaction between sex and genotype for state 3G with increased respiration in male *APOE3* mice and no changes observed in state 3S respiration (**Figure 2C, D**). Uncoupled carbohydrate driven respiration also showed a main effect of genotype with male *APOE3* mice have increased respiration when compared to *APOE4* males (**Figure 2E**).

**Figure 2.**
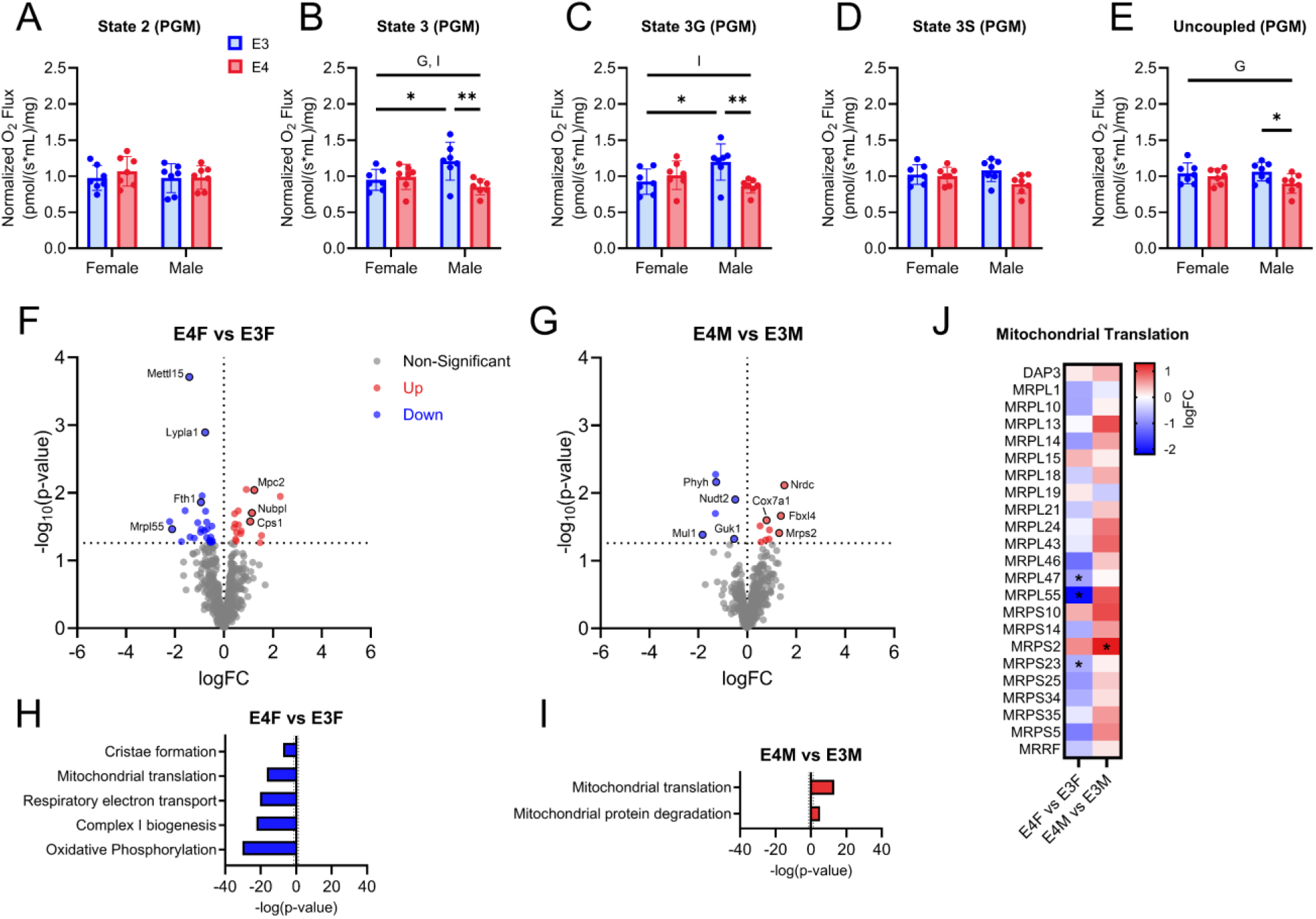
Mitochondrial respiration and isolated brain mitochondria proteomics in male and female *APOE3* and *APOE4* mice. Carbohydrate (PM) driven respiration for A. state 2 B. state 3 C. state 3G D. state 3S and E. uncoupled respiration states. n=7 per genotype and sex. All data are shown as mean ±SD. *p<0.05, ** p<0.01. G=main effect of genotype, I=interaction between genotype and sex. F. Volcano plot showing up and down regulated mitochondrial proteins between female *APOE4* and *APOE3* mice. G. Volcano plot showing up and down regulated proteins between male *APOE4* and *APOE3* mice. H. IPA showing up and down regulated pathways between female *APOE4* and *APOE3* mice. I. IPA showing up and down regulated pathways between male *APOE4* and *APOE3* mice. J. Heatmap of proteins involved in mitochondrial translation between *APOE4* and *APOE3* mice.

We next examined isolated mitochondrial proteomics from *APOE* TR mouse brain samples. After LC/MS analysis of samples, we cross referenced our protein lists with MitoCarta3.0. Female *APOE4* mice had increased expression of mitochondrial pyruvate carrier (Mpc2), complex I assembly (Nubpl), and urea cycle (Cps1) proteins (**Figure 2F**). Mitochondrial ribosome (Mrpl55), RNA methylation (Mett15), lysophospholipase activity (Lypla1), and iron homeostasis (Fth1) proteins were down regulated in female *APOE4* mice (**Figure 2F**). Male *APOE4* mice had increased expression of metalloendopeptidase (Nrdc), cytochrome oxidase (Cox7a1), mitochondrial DNA integrity/mitophagy inhibition (Fbxl14), and mitochondrial ribosome (Mrps2) proteins (**Figure 2G**). Proteins involved in mitochondrial DNA maintenance/GDP formation (Guk1), α-oxidation/lipid metabolism (Phys), ubiquitin ligase (Mul1), and viral RNA degradation/AMP metabolism (Nudt2) were down regulated in male *APOE4* mice when compared to *APOE3* males (**Figure 2G**).

IPA of isolated brain mitochondrial proteomics showed a down regulation of mitochondrial translation, cristae formation, respiratory electron transport, complex I biogenesis, and oxidative phosphorylation in *APOE4* female mice (**Figure 2H**). Male *APOE4* mice showed an upregulation of mitochondrial translation and mitochondrial protein degradation pathways (**Figure 2I**). Interestingly, sex specific differences were observed in mitochondrial translation, where *APOE4* females had a down regulation of this pathway while it was upregulated in male *APOE4* mice (**Figure 2H-J**). Specific mitochondrial ribosomal proteins including MRPL47 and MRPS23 were down regulated in female *APOE4* mice while MRSP2 was upregulated in male *APOE4* mice (**Figure 2J**). Sex differences were also observed in isolated brain mitochondria (**Supplemental Figure 2**).

### Proteomics reveals APOE genotype dependent shifts in iAstrocytes and iNeurons

To examine cell type-specific effects of *APOE* genotype, we generated induced pluripotent stem cell derived neurons (iNeurons) and astrocytes (iAstrocytes) from two isogenic pairs of iPSCs and measured proteomic (Pair A only) and mitochondrial/bioenergetic outcomes (**Figure 3A**). Characterization of iAstrocytes revealed expression of the astrocytic marker S100B, while iNeurons expressed the neuronal marker MAP2 (**Figure 3B**). APOE protein expression in *APOE4* iAstrocytes was decreased when compared to *APOE3* cells, an effect not observed in iNeurons, which showed no difference (**Figure 3C, D**). Surprisingly, *APOE4* iAstrocytes showed an upregulation of 2344 proteins (33.40%) and a down regulation of 2463 proteins (35.09%) (**Figure 3E**). Specific proteins of interest that were upregulated in *APOE4* iAstrocytes included those associated with inflammation (CD38, LXN, ANXA3), redox (ISLR), retinoic acid metabolism (CRABP1), cell stability/movement and tight junctions (ANK1, TJP2/ZO-2) (**Figure 3E**). Proteins that were down regulated in *APOE4* iAstrocytes included those associated with axon guidance/VEGF receptor ligand (NRP2), protein trafficking (SORT1), synaptogenesis (NCAM1), chaperone/lipid transport (CLU) and ER membrane trafficking and ER stress (RTN1) (**Figure 3E**).

**Figure 3.**
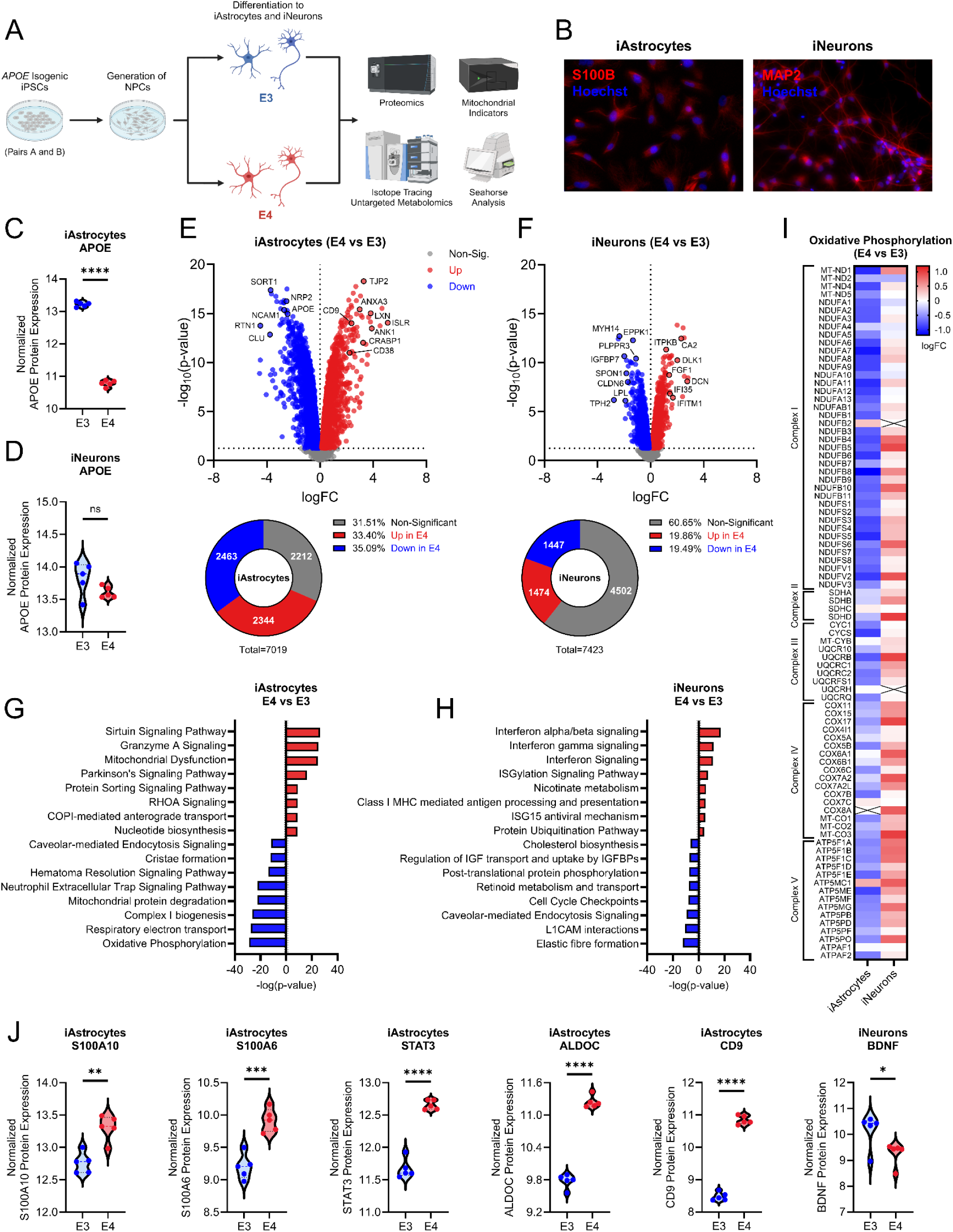
Proteomic analysis of isogenic *APOE* iAstrocytes and iNeurons. A. Study schematic. Two pairs of isogenic iPSCs with either *APOE3* or *APOE4* genotype were used to generate iAstrocytes and iNeurons and examine proteomic and bioenergetic outcomes. B. S100B (red) and Hoechst (blue) staining in iAstrocytes, MAP2 (red) and Hoechst (blue) staining in iNeurons. C. APOE protein expression in iAstrocytes and D. APOE protein expression in iNeurons. E. Volcano plot showing up and down regulated proteins in iAstrocytes between *APOE4* and *APOE3*. F. Volcano plot showing up and down regulated proteins in iNeurons between *APOE4* and *APOE3*. G. IPA showing up and down regulated pathways in iAstrocytes between *APOE4* and *APOE3*. H. IPA showing up and down regulated pathways in iNeurons between *APOE4* and *APOE3*. I. Heatmap of the proteins involved in oxidative phosphorylation comparing *APOE* genotype mediated effects between iAstrocytes and iNeurons. n=5 per group. J. Violin plots showing disease associed markers S100A10, S100A6, STAT3, ALDOC, and CD9 in iAstrocytes along with BDNF levels in iNeurons. ns=not significant, *p<0.05, **p<0.01, ***p<0.001, ****p<0.0001.

iNeurons, however, had far fewer differentially expressed proteins than iAstrocytes, with 1,474 upregulated (19.86%) and 1,447 downregulated (19.49%) proteins in *APOE4* derived cells when compared to *APOE3* (**Figure 3F**). Selected proteins that were upregulated in *APOE4* iNeurons included those associated with inflammation (IFI35, IFITMI), extracellular matrix (DCN), calcium signaling (ITPKB), neuronal signaling/metalloenzyme (CA2) and axon regeneration/neurotrophic factors (DLK1, FGF1) (**Figure 3F**). Proteins that were down regulated in *APOE4* iNeurons included those associated with axon guidance (PLPPR3, SPON1), actin/cytoskeleton organization and cell barrier (MYH14, EPPK1, CLDN6), lipid metabolism (LPL), serotonin synthesis (TPH2), and insulin like growth factor signaling (IGFBP7) (**Figure 3F**).

IPA showed an upregulation of sirtuin signaling, granzyme A signaling, mitochondrial dysfunction, protein sorting, RHOA signaling, COPI anterograde transport, Parkinson’s signaling, and nucleotide biosynthesis pathways in *APOE4* iAstrocytes (**Figure 3G**). Down regulated pathways in *APOE* iAstrocytes included endocytosis, cristae formation, neutrophil extracellular trap signaling, mitochondrial protein degradation, complex I biogenesis, respiratory electron transport/oxidative phosphorylation, and hematoma resolution signaling (**Figure 3G**). In iNeurons, IPA showed an upregulation of interferon signaling, ISGylation, nicotinate metabolism, class I MHC antigen processing/presentation, ISG15 antiviral, and ubiquitination pathways in *APOE4* cells (**Figure 3H**). Down regulated pathways in *APOE4* iNeurons included cholesterol biosynthesis, IGF transport/uptake, protein phosphorylation, retinoid metabolism, cell cycle check points, endocytosis, L1CAM interactions, and elastic fibre formation (**Figure 3H**). Interestingly, oxidative phosphorylation proteins were largely down regulated in *APOE4* iAstrocytes while upregulated in *APOE4* iNeurons (**Figure 3I**). Disease associated astrocytic (DAA) markers were upregulated in *APOE4* iAstrocytes, including S100A10, S100A6, STAT3, ALDOC, and CD9 (**Figure 3J**). Brain derived neurotrophic factor (BDNF) was down regulated in *APOE4* iNeurons (**Figure 3J**).

### Mitochondrial function in iAstrocytes and iNeurons is altered by APOE genetic variation

We next examined mitochondrial function in both isogenic pairs of derived iAstrocytes. *APOE4* iAstrocytes derived from both Pair A and B had reduced basal, maximal, proton leak, and ATP-production linked respiration (**Figure 4A-D**). *APOE4* iAstrocytes also showed reduced complex I, II, and IV flux/driven respiration in both isogenic pairs, with disparate results for complex III (reduced in Pair A) (**Figure 4E, F**). Mitochondrial membrane potential, mitochondrial superoxide production, hydrogen peroxide (H_2_O_2_), intracellular calcium (Ca^2+^), and mitochondrial Ca^2+^ were increased in both Pair A and B derived *APOE4* iAstrocytes (**Figure 4G, H**). Isogenic *APOE4* iNeurons derived from both Pair A and B had reduced maximal respiration (**Figure 5A-D**). Disparate results were observed for basal respiration and proton leak which were reduced in group B only (**Figure 5A-D**). Pair B *APOE4* iNeurons from showed reduced complex I, II, and III flux which was not observed in group A (**Figure 5E, F**). Both Pair A and B *APOE4* iNeurons had increased complex IV flux/driven respiration (**Figure 5E, F**). Mitochondrial membrane potential, H_2_O_2_, and total cellular calcium were increased in Pair B *APOE4* iNeurons but not Pair A (**Figure 5G, H**). Mitochondrial superoxide was increased in Pair A *APOE4* iNeurons with an increased trend in Pair B (**Figure 5G, H**). Mitochondrial Ca^2+^ levels were increased in both Pair A and B derived *APOE4* iNeurons (**Figure 5G, H**).

**Figure 4.**
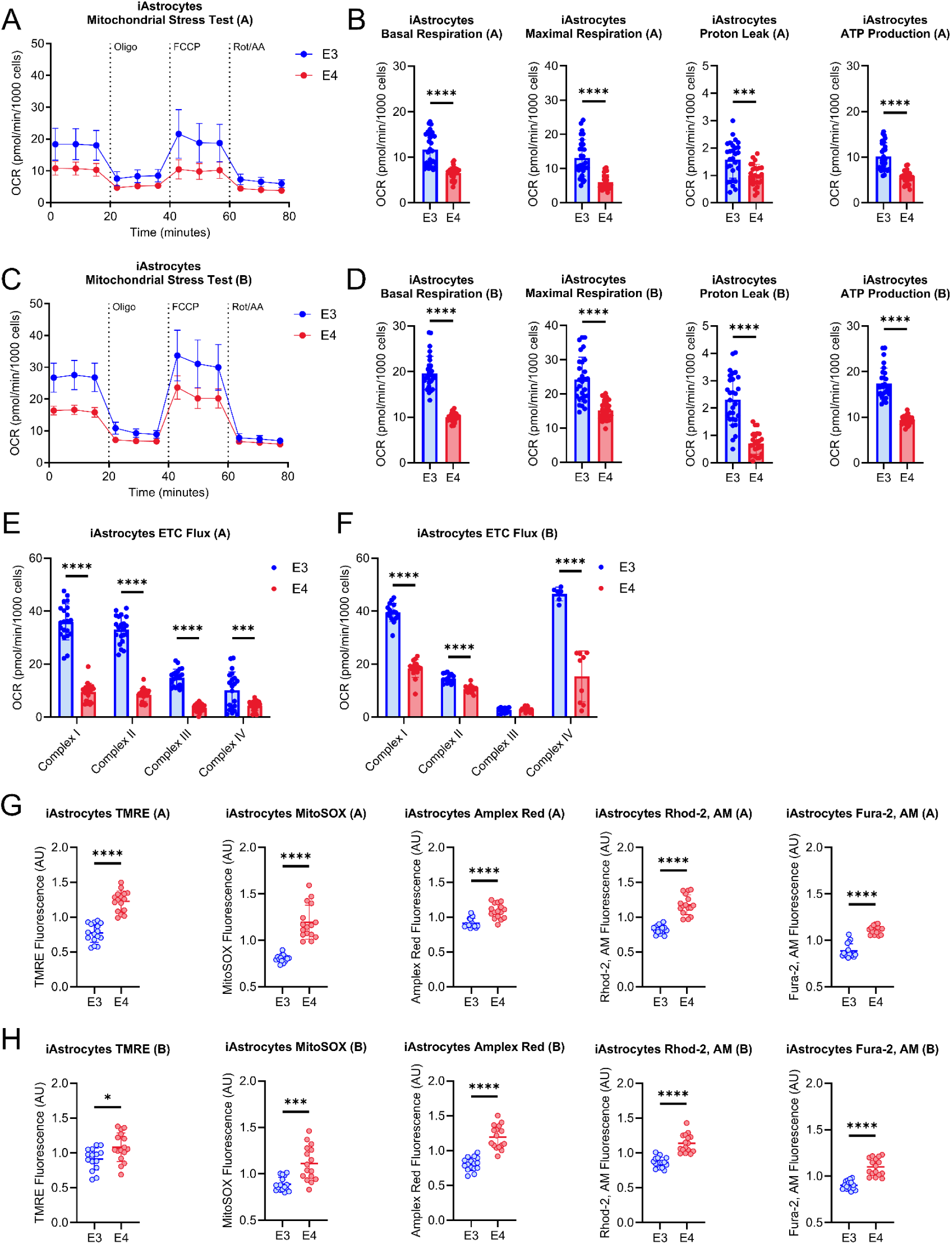
Mitochondrial function in iAstrocytes. A. Mitochondrial stress test (MST) tracing from isogenic Pair A. B. Quantification of basal respiration, maximal respiration, proton leak, and ATP-production from isogenic Pair A. C. MST tracing from isogenic Pair B. D. Quantification of basal respiration, maximal respiration, proton leak, and ATP-production from isogenic Pair B. E. Electron transport chain (ETC) flux/driven respiration through complex I, II, III, and IV in isogenic Pair A. F. ETC flux/driven respiration through complex I, II, III, and IV in isogenic Pair B. G. TMRE, MitoSOX, Amplex Red, Rhod-2, AM, and Fura-2, AM fluorescence intensity from isogenic Pair A. H. TMRE, MitoSOX, Amplex Red, Rhod-2, AM, and Fura-2, AM fluorescence intensity from isogenic Pair B. n=16. All data are shown as mean ±SD. ns=not significant, *p<0.05, **p<0.01, ***p<0.001, ****p<0.0001.

**Figure 5.**
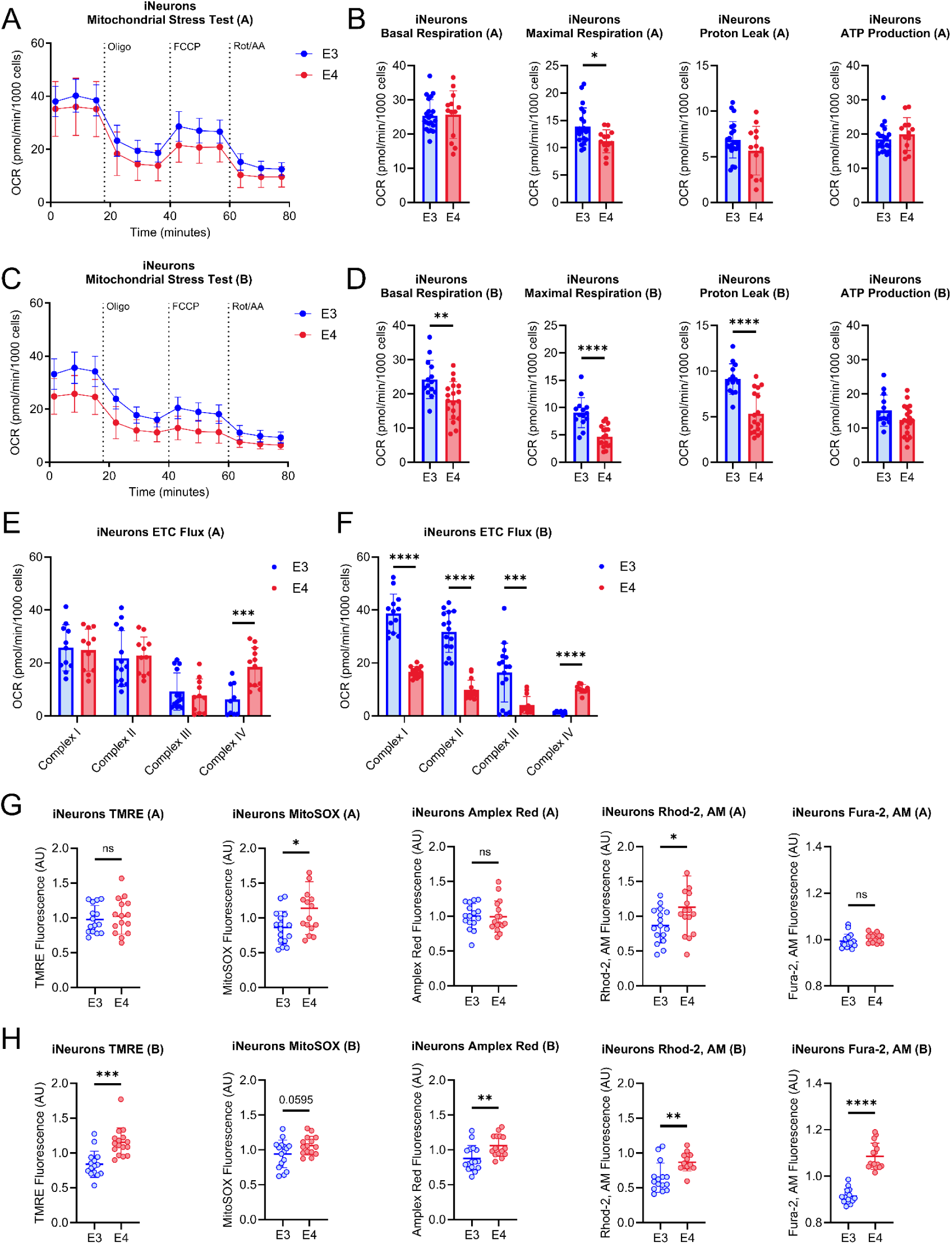
Mitochondrial function in iNeurons. A. Mitochondrial stress test (MST) tracing from isogenic Pair A. B. Quantification of basal respiration, maximal respiration, proton leak, and ATP-production from isogenic Pair A. C. MST tracing from isogenic Pair B. D. Quantification of basal respiration, maximal respiration, proton leak, and ATP-production from isogenic Pair B. E. Electron transport chain (ETC) flux/driven respiration through complex I, II, III, and IV in isogenic Pair A. F. ETC flux/driven respiration through complex I, II, III, and IV in isogenic Pair B. G. TMRE, MitoSOX, Amplex Red, Rhod-2, AM, and Fura-2, AM fluorescence intensity from isogenic Pair A. H. TMRE, MitoSOX, Amplex Red, Rhod-2, AM, and Fura-2, AM fluorescence intensity from isogenic Pair B. n=16. All data are shown as mean ±SD. ns=not significant, *p<0.05, **p<0.01, ***p<0.001, ****p<0.0001.

### APOE4 increases glycolysis in iAstrocytes and iNeurons

Glycolytic flux was increased in *APOE4* iAstrocytes, including glycolytic capacity, glycolytic reserve, and a trend for increased non-glycolytic extracellular acidification rate (**Figure 6A, B**). Glycolytic flux was also increased in *APOE4* iNeurons including basal glycolysis, glycolytic capacity, glycolytic reserve, and non-glycolytic extracellular acidification rates (**Figure 6C, D**). To further interrogate these findings, we measured fuel source dependency and capacity for use in mitochondrial respiration. Glucose and fatty acid fuel source dependency and capacity were both increased in *APOE4* iAstrocytes (**Figure 6E, F**). iNeurons had decreased glucose and glutamine fuel dependency with increased capacity to use glucose and decreased capacity to use fatty acids as fuel sources (**Figure 6G, H**). Furthermore, glycolytic pathway protein expression was largely increased in both iAstrocytes and iNeurons (**Figure 6I**).

**Figure 6.**
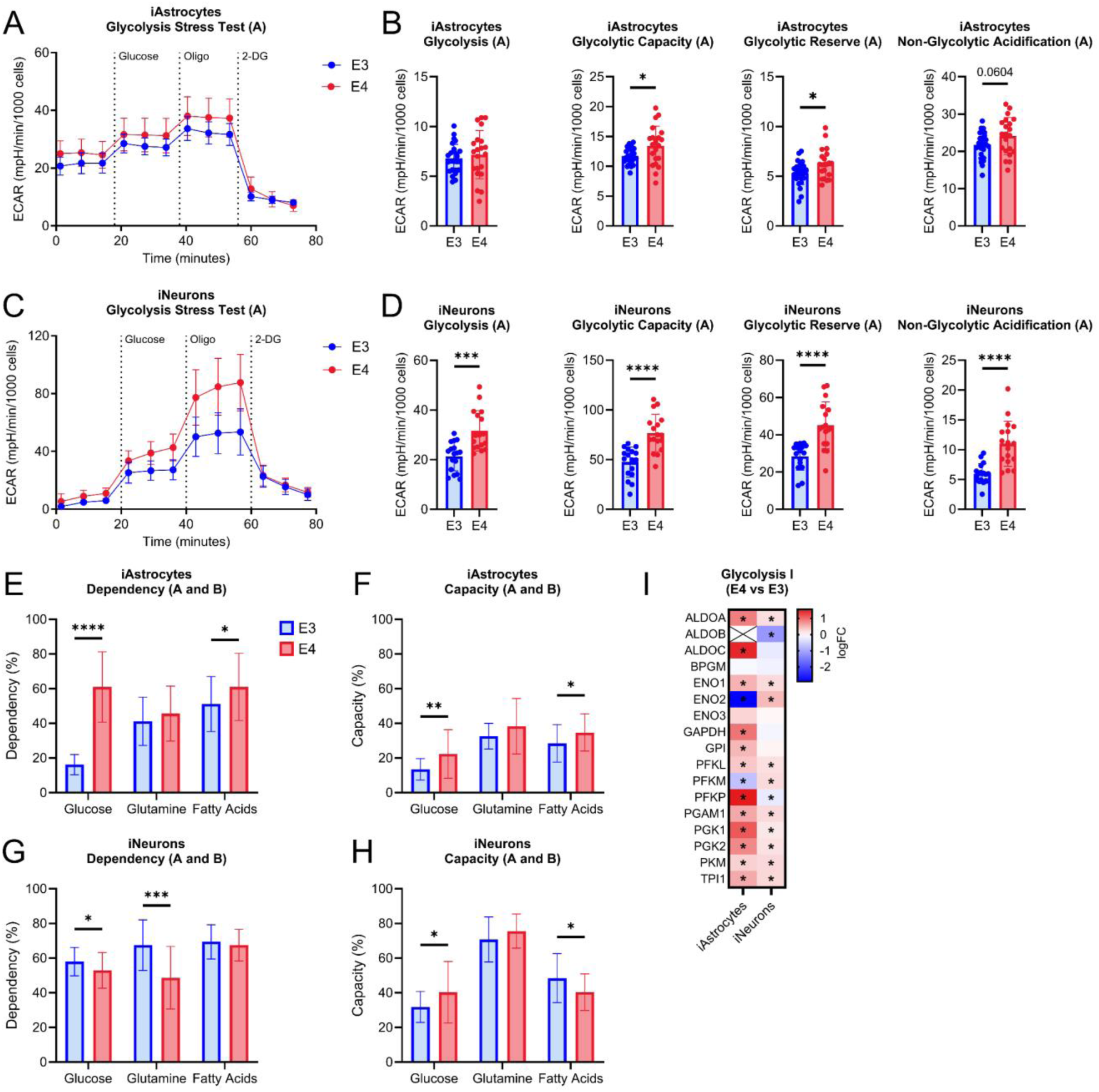
Glycolytic function in iAstrocytes and iNeurons. A. Glycolytic stress test (GST) tracing from Pair A iAstrocytes. B. Quantification of basal glycolysis, glycolytic capacity, glycolytic reserve, and non-glycolytic acidification from iAstrocytes. C. GST tracing from Pair A iNeurons. D. Quantification of basal glycolysis, glycolytic capacity, glycolytic reserve, and non-glycolytic acidification from iNeurons. E. Glucose, glutamine, and fatty acid fuel dependency in Pair A and B iAstrocytes. F. Glucose, glutamine, and fatty acid fuel capacity in Pair A and B iAstrocytes. G. Glucose, glutamine, and fatty acid fuel dependency in Pair A and B iNeurons. H. Glucose, glutamine, and fatty acid fuel capacity in Pair A and B iNeurons. I. Heatmap of proteins involved in glycolysis I network comparing iAstrocyte and iNeuron outcomes between *APOE4* and *APOE3*. All data are shown as mean ±SD. *p<0.05, **p<0.01, ***p<0.001, ****p<0.0001.

### 13C-Glucose labeling of metabolites is altered by APOE genotype in iAstrocytes and iNeurons

To further understand glucose metabolic pathways that differ based on *APOE* genotype, we employed stable isotope labeling in iAstrocytes and iNeurons using [U-^13^C_6_]-glucose (**Figure 7A**). We first examined effects on TCA cycle metabolites and intermediates. *APOE4* iAstrocytes had increased labeling of aspartic acid for M+1, M+2, and M+4 isotopologues (**Figure 7B**). No differences were observed for cis-aconitic acid labeling in iAstrocytes (**Figure 7C**). Additionally, there was increased labeling of the M+1, M+2, M+4, and M+5 isotopologues for glutamic acid but decreased labeling of the M+3 isotopologue in *APOE4* iAstrocytes (**Figure 7D**). Unlabeled succinate was significantly increased in *APOE4* iAstrocytes with reduced labeling for the M+2 isotopologue (**Figure 7E**). Interestingly, there was both increased and decreased labeling of malic acid (**Figure 7F**). We specifically observed a decrease for the M+3 isotopologue and an increase in the M+4 isotopologue in *APOE4* iAstrocytes.

**Figure 7.**
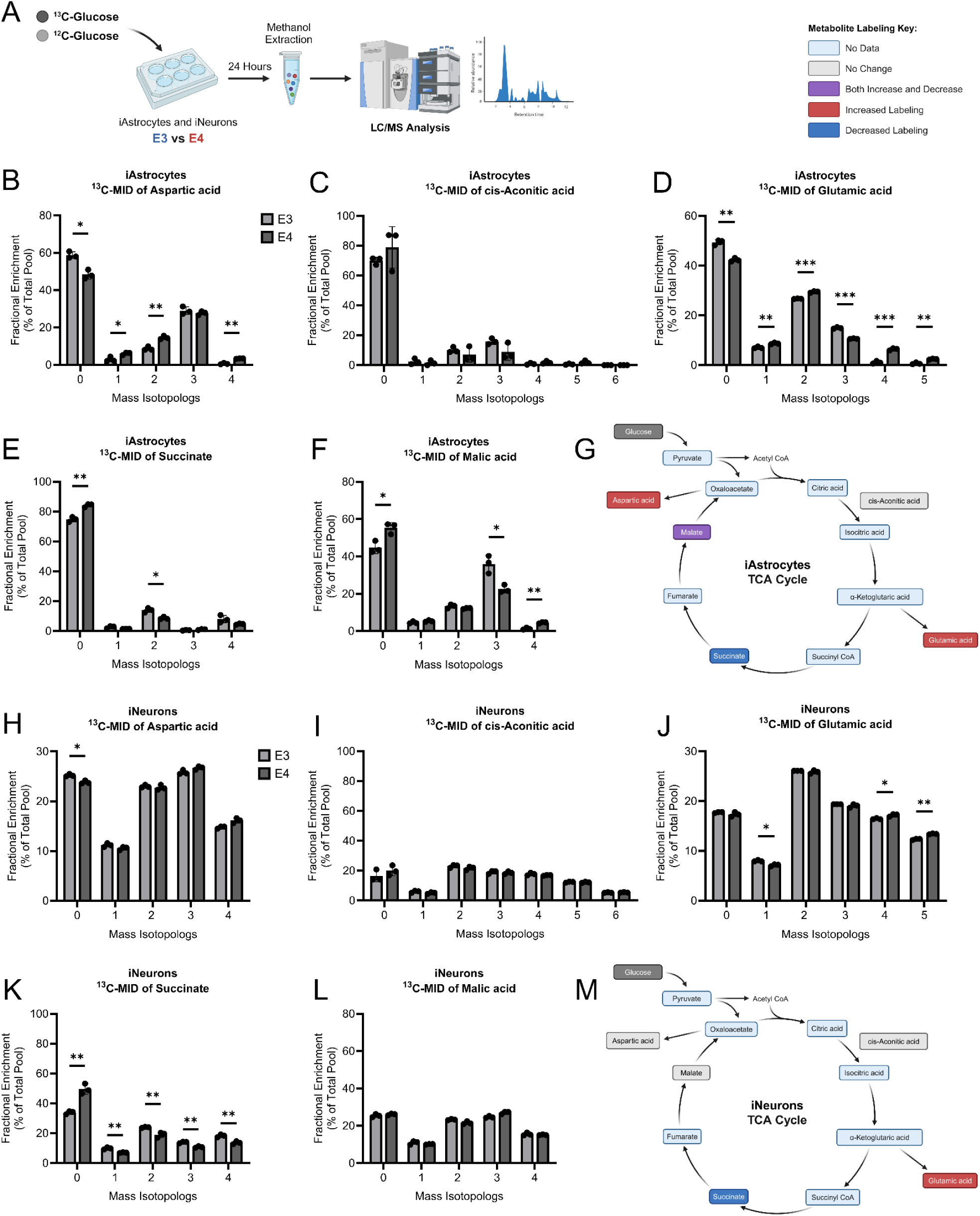
^13^C-Glucose stable isotope labeling of TCA cycle metabolites and intermediates in iAstrocytes and iNeurons. A. Schematic of stable isotope labeling experiment using ^13^C-glucose. iAstrocyte mass isotopologue labeling of B. aspartic acid, C. cis-aconitic acid, D. glutamic Acid, E. succinate, F. malic acid. G. Overview of TCA cycle metabolite labeling outcomes in iAstrocytes. iNeuron mass isotopologue labeling of H. aspartic acid, I. cis-aconitic acid, J. glutamic acid, K. succinate, L. malic acid. M. Overview of TCA cycle metabolite labeling outcomes in iNeurons. n=3 per group. All data are shown as mean ±SD. *p<0.05, **p<0.01, ***p<0.001. Data were analyzed using a multiple un-paired t-test with Holm-Sidak correction.

*APOE4* iNeurons had decreased unlabeled aspartic acid and no differences were observed for cis-aconitic acid labeling (**Figure 7H, I**). We also observed that *APOE4* iNeurons had increased labeling with M+4 and M+5 isotopologues for glutamic acid but decreased labeling of the M+1 isotopologue (**Figure 7J**). Unlabeled succinate was increased in *APOE4* iNeurons with reduced labeling of the M+1, M+2, M+3, and M+4 isotopologues (**Figure 7K**). Malic acid labeling in iNeurons showed no differences with *APOE* genotype (**Figure 7L**). These data suggest increased labeling of glutamic acid and decreased labeling of succinate with respect to glucose metabolism in *APOE4* iNeurons.

Due to differences observed in ^13^C-glucose labeling of TCA cycle metabolites— specifically malate, aspartate, and glutamate—in *APOE4* iAstrocytes, we further examined proteins associated with their metabolism. Analysis of TCA cycle proteins revealed an overall decrease in *APOE4* iAstrocytes, while iNeurons exhibited minimal changes, with a few proteins upregulated in *APOE4* cells (**Figure 8A**). Given these changes to TCA cycle proteins and altered ^13^C-glucose labeling, we next investigated proteins associated with the malate-aspartate shuttle in iAstrocytes. As stated earlier, *APOE4* iAstrocytes exhibit increased M+4 malate; however, protein levels of pyruvate dehydrogenase subunits PDHA1 and PDHB were decreased, suggesting impaired glucose carbon entry into the TCA cycle (**Figure 7F and Figure 8B, C**). Interestingly, there was an upregulation fumarate hydratase (FH) which is responsible for converting fumarate to malate (**Figure 8D**). We also observed an increase in cytoplasmic malate dehydrogenase (MDH1) and a decrease in mitochondrial malate dehydrogenase (MDH2), which are responsible for the reversible conversion of malate to oxaloacetate (**Figure 8E, F**). However, increased pyruvate carboxylase (PC) alongside decreased M+3 malate suggests that while anaplerotic capacity/activity is elevated, its contribution to TCA cycle intermediates may be limited (**Figure 7F and Figure 8G**). In combination with increased levels of PC, we also observe increased levels of cytoplasmic aspartate aminotransferase (GOT1) and a decrease in mitochondrial aspartate aminotransferase (GOT2), which catalyzes the conversion of oxaloacetate to aspartic acid (**Figure 8H, I**). Notably, MDH1 and GOT1 protein levels were positively correlated in *APOE4* iAstrocytes, a relationship absent in *APOE3* cells (**Figure 8J**). No significant correlations were observed between MDH2 and GOT2 levels in either genotype (**Figure 8K**). To further support altered malate-aspartate shuttle activity, we investigated levels of mitochondrial transporters SLC25A11 and SLC25A13 in *APOE4* iAstrocytes, which are critical for shuttling malate and aspartate across the mitochondrial membrane and found that both were significantly decreased. (**Figure 8L, M**). Collectively, these results support a metabolic shift favoring the cytosolic conversion of glucose-derived pyruvate to oxaloacetate and aspartic acid, bypassing the TCA cycle.

**Figure 8.**
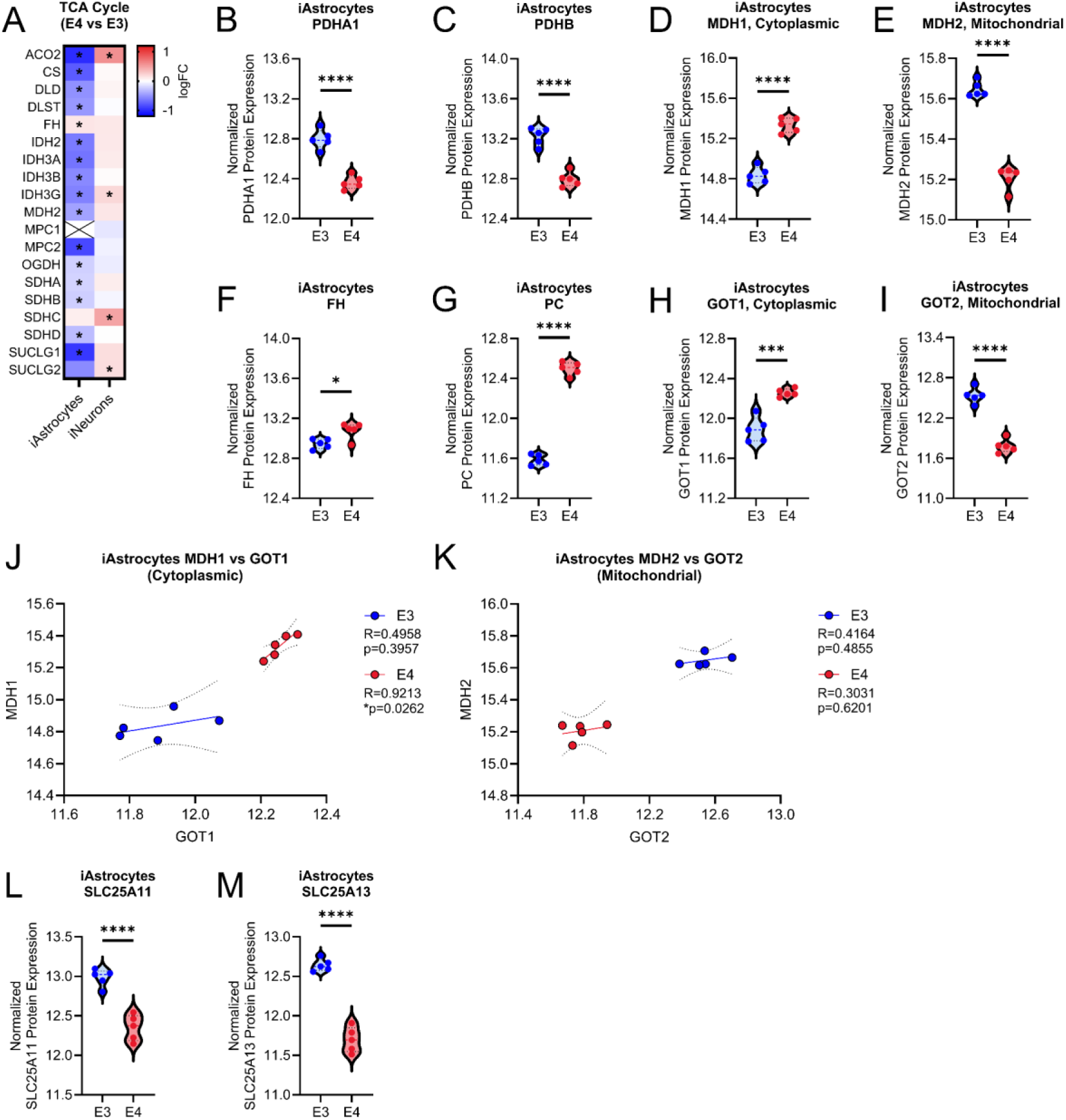
Proteins involved in the TCA cycle and malate-aspartate shuttle in isogenic *APOE* iAstrocytes. A. Heatmap of TCA cycle proteins in iAstrocytes and iNeurons. Violin plots showing levels of TCA cycle and related proteins in iAstrocytes including B. PDHA1, C. PDHB, D. FH, E. MDH1, F. MDH2, G. PC, H. GOT1, I. GOT2. J. Correlation analysis of MDH1 and GOT1 protein expression levels. K. Correlation analysis of MDH2 and GOT2 protein expression levels. Violin plots showing levels of L. SLC25A11 and M. SLC25A13. *p<0.05, ***p<0.001, ****p<0.0001.

Next, we sought to look at nucleotide, energy transfer, and redox pair molecules. *APOE4* iAstrocytes and iNeurons had increased ^13^C-glucose labeling of ATP (**Figure 9A**). Additionally, there was increased ^13^C-glucose labeling of ADP, GDP, NADP, NAD, NADH, UTP, UDP, and UDP-GlcNAc in *APOE4* iAstrocytes (**Figure 9B, D-J**). *APOE4* iNeurons had increased ^13^C-glucose labeling of GTP, NAD, NADH, UTP, UDP, and UDP-N-acetylglucosamine (UDP-GlcNAc) (**Figure 9C, F-J**). Due to an increase in ^13^C-glucose labeled UDP-GlcNAc, we analyzed proteins in the hexosamine and GlcNAc synthesis pathway which had increased expression in both *APOE4* iAstrocytes and iNeurons (**Figure 9K**). Levels of O-GlcNAc were also higher in *APOE4* cells (**Figure 9L**). The hexosamine biosynthetic pathway and GlcNAc synthesis pathway are outlined in **Figure 9M**, with an emphasis on metabolites and proteins with altered expression profiles in *APOE4* derived cells. Overall, this data suggests increased flux of ^13^C-glucose through this pathway. Additional metabolites were measured in **Supplemental Figure 3**. This includes a decrease in the amount of ^13^C-glucose labeled fructose 1,6-bisphosphate and N-acetyl cysteine (NAC) in APOE4 iNeurons (**Supplemental Figure 3A, E**). However, no changes were observed for glycerol 3-phosphate, lactate, and alanine (**Supplemental Figure 3B-D**).

**Figure 9.**
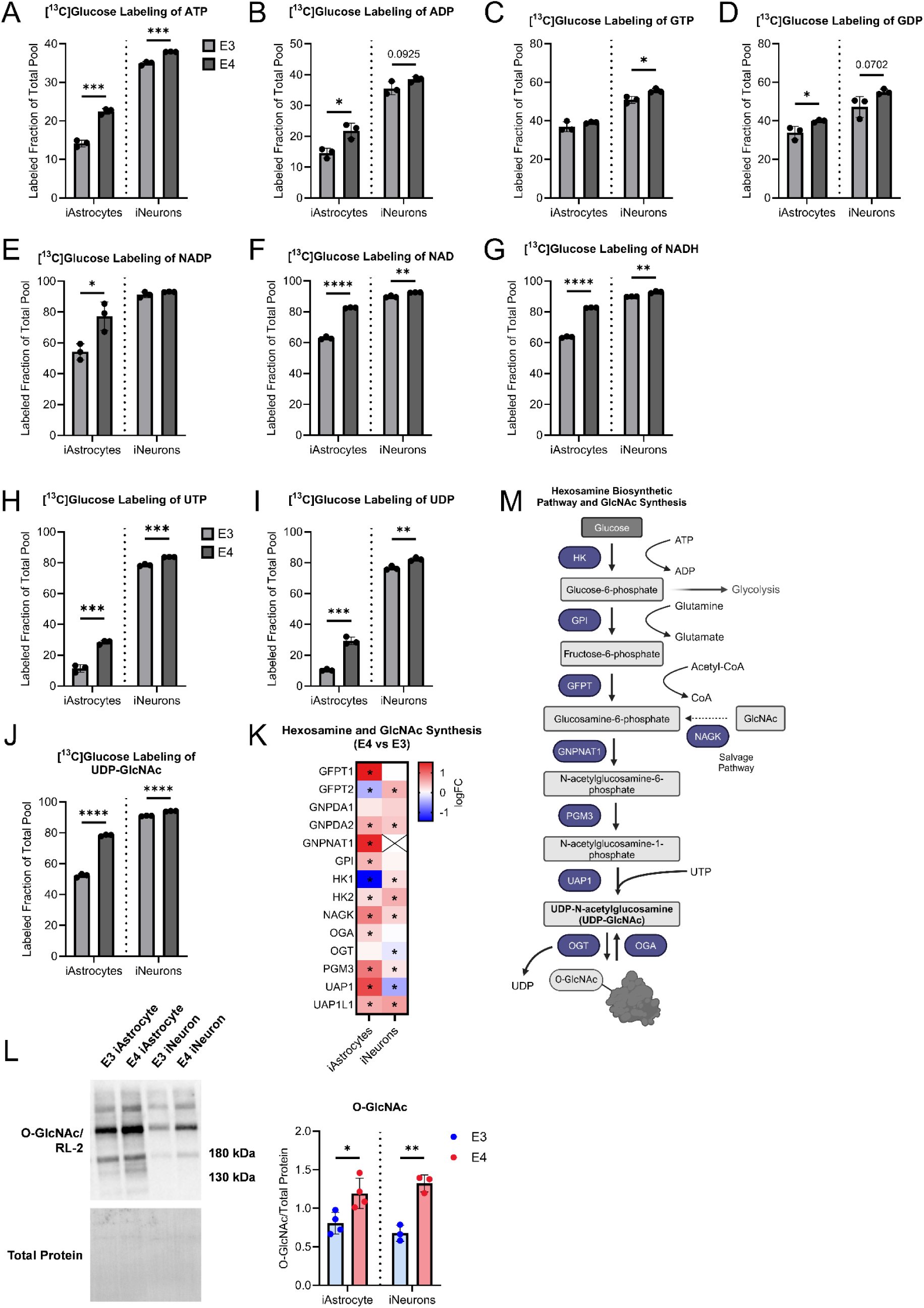
Stable isotope labeling of nucleotide/energy transfer/redox pair molecules, and evaluation of hexosamine/GlcNAc pathway in iAstrocytes and iNeurons. iAstrocyte and iNeuron labeled fraction of total pool for A. ATP, B. ADP, C. GTP, D. GDP, E. NADP, F. NAD, G. NADH, H. UTP, I. UDP, J. UDP-GlcNAc. K. Heatmap of proteins involved in the Hexosamine and GlcNAc synthesis pathway in *APOE4* vs. *APOE3* iAstrocytes and iNeurons. L. O-GlcNAc western blot on *APOE3* and *APOE4* iAstrocytes and iNeurons with densitometry quantification. M. Schematic overview of hexosamine and GlcNAc synthesis pathway. n=3 per group. All data are shown as mean ±SD. *p<0.05, **p<0.01, ***p<0.001, ****p<0.0001.

## Discussion

The overall goal of this study was to examine the effects of *APOE* genotype on brain health, metabolism, and mitochondrial function. To this end, we examined whole brain and isolated brain mitochondrial proteomics coupled with mitochondrial function in *APOE* TR mice harboring *APOE3* or *APOE4* genes. We complimented mouse data by examining cell specific effects of *APOE* genotype using isogenic iPSC-derived astrocytes and neurons. These data revealed reduced APOE protein expression in *APOE4* mouse brain and iAstrocytes, but not iNeurons. Observations of reduced APOE levels have been previously reported by other groups (29, 31). While a definitive conclusion cannot be made with current data, it is likely that whole brain reduced APOE protein expression in *APOE4* models is due to astrocytic or glial cell phenotypes. The significance of reduced *APOE4* expression includes dysfunctional lipid trafficking and/or reduced lipid metabolism.

Whole brain proteomic data indeed revealed significant bioenergetic and lipid metabolism changes. Some mitochondrial proteins were increased, which participate in electron transport chain function at complex IV (mt-Co3) and complex II (Sdhd) in female *APOE4* mice, but they also showed reduced expression of a key ketone metabolism protein, Hmgcs2. Pathway analysis revealed reduced signaling through metabolic pathways as well, including mitochondrial translation and response to amino acid deficiency. Upregulated pathways revealed pathways associated with autophagy (neddylation) and intracellular vesicle trafficking (RAB geranylgeranylation) suggesting activation of compensatory mechanisms in response to metabolic stress. Decreased expression of ketone metabolism enzyme Hmgcs2 with increased expression of complex II component Sdhd suggests a concerted effort to overcome impaired lipid metabolism.

Male *APOE4* mice had similar overall responses to female mice, with regards to metabolic stress and alterations to lipid metabolism. One key difference was increased inflammatory associated protein expression with Cfb and Gsdma being upregulated. Interestingly, Cfb has been previously linked to neurodegeneration (49). Male *APOE4* mice also had reduced activation of the mitochondrial dysfunction pathway with increased activation of cristae formation, suggesting an effort to compensate for metabolic stress or reduced mitochondrial mass. Mitochondrial function from isolated brain mitochondria agrees with our proteomic data suggesting compensatory efforts to overcome metabolic dysfunction. Male *APOE4* mice had decreased mitochondrial respiration for state 3 with pyruvate, malate, and glutamate as substrates. State 3 respiration is a measure of electron transport chain function, as substrates and ADP are present at saturating levels. Synaptic pathways were also activated which largely revealed increased inhibitory transmitter signaling. Additionally, upregulation of synaptogenesis suggests a compensatory effect in *APOE4* mice as previous work has found the *APOE4* allele is associated with altered synaptic density (50–52).

Proteomic analysis of isolated mitochondria from brain further revealed a down regulation of lipid metabolism pathway components (Phyh in males, Lypla1 in females) with an upregulation of certain electron transport chain components (Nubpl in females, Cox7a1 in males). These changes in protein expression are interesting since we see a downregulation of pathways involved in complex I biogenesis, respiratory chain/oxidative phosphorylation, and mitochondrial translation. Further studies are needed to elucidate mechanisms of these outcomes, with specific experiments to understand if these changes are compensatory due to a response from metabolic stress.

When looking at proteomic differences in iAstrocytes and iNeurons we found a greater number of DE proteins in iAstrocytes than iNeurons. This is an interesting finding since astrocytes are known to be the primary producers of APOE in the CNS. *APOE4* derived iAstrocytes showed increased inflammatory protein expression including those associated with DAA phenotypes in AD (53–56). Using IPA we found that the mitochondrial dysfunction pathway was upregulated with cristae formation, complex I biogenesis, and respiratory chain/oxidative phosphorylation being downregulated in *APOE4* derived iAstrocytes. *APOE4* derived iNeurons also increased inflammatory protein expression and calcium signaling with decreased expression of proteins involved in lipid metabolism (specifically CLU and SORT1). Interferon signaling was significantly upregulated in iNeurons with *APOE4* in addition to other antiviral inflammatory proteins. We also found class I MHC mediated antigen processing and presentation being upregulated in *APOE4* iNeurons which is consistent with a previous study (36). Cholesterol biosynthesis, insulin like growth factor signaling were all downregulated in *APOE4* iNeurons. Despite both cell types showing profound metabolic stress and mitochondrial dysfunction, oxidative phosphorylation associated proteins were downregulated in *APOE4* iAstrocytes and upregulated in *APOE4* iNeurons. These results suggest cell type specific responses to stress and could reflect important considerations for therapeutic development.

Mitochondrial function in iAstrocytes and iNeurons revealed reduced mitochondrial respiration and electron transport chain function in *APOE4* derived cells with increased indices of oxidative stress and mitochondrial calcium. While there were some disparities between isogenic pairs, the data overall suggest *APOE4* induces mitochondrial dysfunction and metabolic stress in iAstrocytes and iNeurons. Reductions in mitochondrial respiration were coupled with increased glycolytic flux, increased dependency/capacity on glucose fuel use in both iAstrocytes and iNeurons with *APOE4*. Alterations to fuel source dependency and capacity can lead to competitive metabolic stress between iAstrocytes and iNeurons. Typically, astrocytes are more glycolytic, and neurons rely more on mitochondrial respiration for energy production. Astrocytes shuttle lactate to neurons to fuel mitochondrial respiration and if this metabolic relationship is disrupted it could negatively impact cellular functions and synaptic transmission. These findings of decreased mitochondrial function and a switch to glycolysis is consistent with previous studies (33, 35). Overall, we found that proteomic and functional differences were greater in *APOE* isogenic iAstrocytes than iNeurons.

Stable isotope tracing with ^13^C glucose revealed *APOE* genotype dependent changes in glucose utilization. We found that both *APOE4* iAstrocytes and iNeurons had increased labeling of glutamic acid and aspartic acid which is consistent with a past study in primary astrocytes (33). However, neurons have not been previously investigated in this context. Malic acid labeling was found to be altered in iAstrocytes but not iNeurons. Given the differential labeling of malic acid, we investigated the expression of proteins involved in both the TCA cycle and malate-aspartate shuttle. Surprisingly, we saw differences in these proteins including a decrease in subunits of PDH and an increase of PC consistent with a past study in iPSC-derived astrocytes (35). This group also observed a decrease in levels of MDH2 and GOT2 which was consistent with our findings. However, they did not report on cytoplasmic MDH1 and GOT1 of which we observed increased expression. This is a critical consideration since these proteins can influence the balance and flux of metabolites in the TCA cycle and malate-aspartate shuttle. Collectively, our findings suggest that *APOE4* suppresses mitochondrial TCA cycle activity in iAstrocytes while promoting cytosolic conversion of pyruvate to aspartate.

We finally looked at labeling of nucleotides and energy transfer molecules. Interestingly, there was increased labeling of ATP from ^13^C-glucose in both *APOE4* iAstrocytes and iNeurons. This data tracks with our functional glycolytic assays and indicates a reliance on glucose for ATP production which is consistent with past studies (30, 31, 57). Other molecules including ADP, GTP, GDP, NAD, NADH, UTP, and UDP also had increased labeling in *APOE4* cells when compared to *APOE3*. Levels of ^13^C-glucose labeled UDP-GlcNAc were higher in *APOE4* iAstrocytes and iNeurons. UDP-GlcNAc participates in numerous cellular functions including nutrient sensing and glycosylation (58). We further examined proteins involved in the hexosamine biosynthetic pathway and GlcNAc synthesis and found levels of these proteins were overall increased. Interestingly, there was an increase in N-acetylglucosamine kinase (NAGK), which participates in salvage synthesis, in both *APOE4* iAstrocytes and iNeurons. Given these findings, we next assessed O-GlcNAcylation, a post-translational modification, and observed an increase in *APOE4* cells. This is particularly interesting since APOE is heavily glycosylated in the CNS and suggests enhanced glycosylation by O-GlcNAcylation in *APOE4* iAstrocytes and iNeurons (59).

Taken together, this study demonstrates significant *APOE* genotype dependent shifts in metabolism. We observed changes in whole brain metabolism and mitochondrial function, including cell type-specific alterations. Male and female *APOE4* mice showed alterations in metabolic proteins/pathways and mitochondrial function. Additionally, while *APOE4* iAstrocytes and iNeurons displayed inflammation and a metabolic shift towards glycolysis, changes in mitochondrial dysfunction were disparate between cell types. Nonetheless, future studies are needed to interrogate underlying mechanisms behind this shift in metabolic function due to *APOE* genetic variation. Specific focus should be placed on understanding if early metabolic and inflammatory stress drive the increased risk of AD in *APOE4* carriers.

## Acknowledgements and Funding

This study was supported by the Margaret “Peg” McLaughlin and Lydia A. Walker Opportunity Fund, the University of Kansas Alzheimer’s Disease Center P30AG072973 (HMW, JKM, and JPT), Kansas Center for Metabolism and Obesity Research Center P20 GM144269 (JPT), R01AG078186 (HMW), R00AG056600 (HMW), R01DK121497 supplement (JPT), R01AG069781 (JPT), VA Merit Review grant 1I01BX002567 (JPT), AG078114 (BAK and CRL), Alzheimer Association Grant 23AARG-1023294 (HMW), T32HD057850 (VC), T32 Brain Health (CL and BK), P20GM103418 (CAG). The content is solely the responsibility of the authors and does not necessarily represent the official views of the National Institutes of Health.

## Author Contributions

Acquisition of Funding (JKM, PCG, JPT, and HMW), Investigation (CRL, CNJ, VC, EF, MB, CAG, CJB, XD, CSM, KPS, PP, PC), Resources and Methodology (PP, PC, CS, JKM, PCG, JPT, and HMW), Supervision (JKM, PGM, JPT, HMW), Writing – Original Draft Preparation (CRL, HMW), Writing – Review & Editing (all authors)

---

**Supplemental Figure 1.**
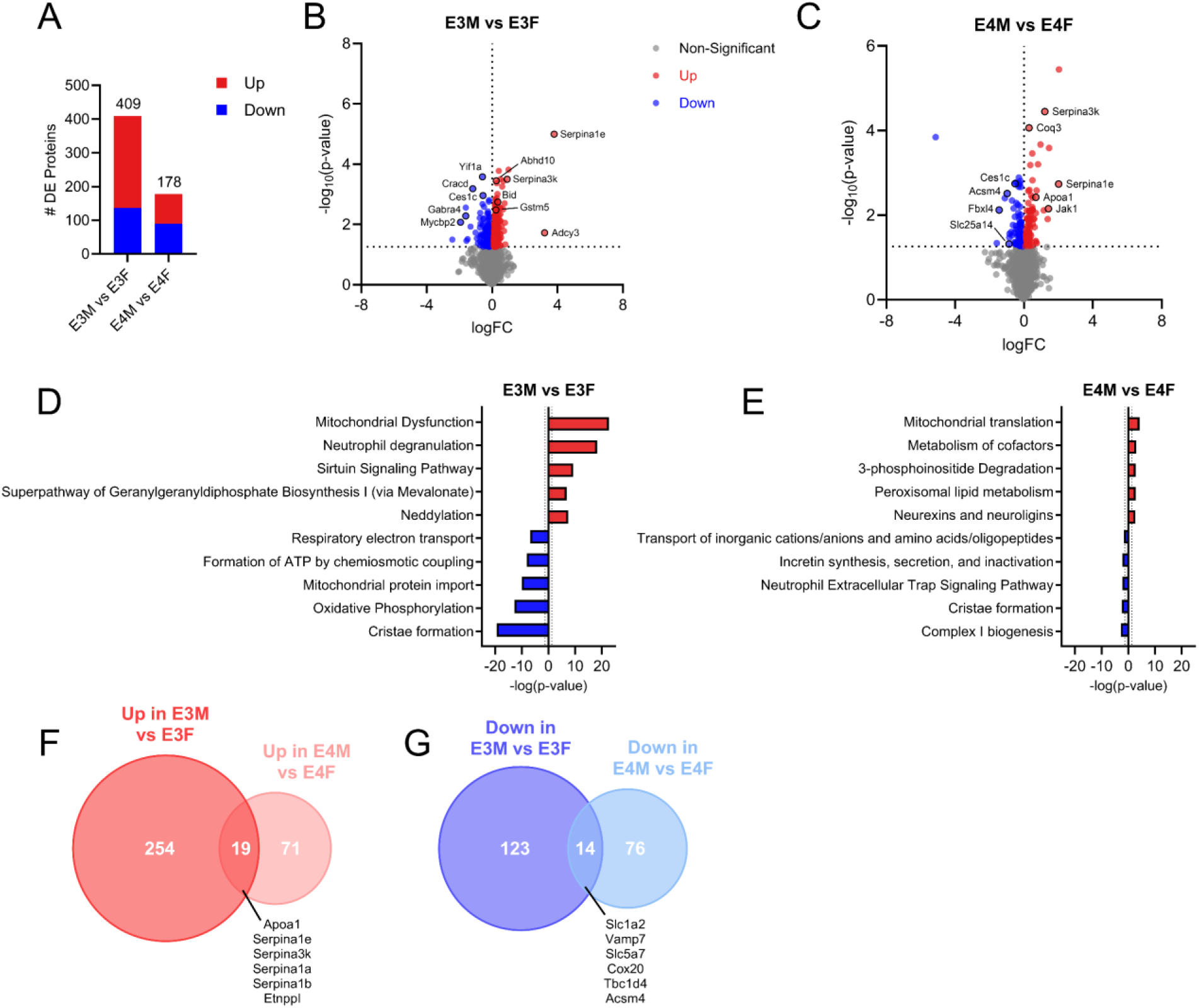
Whole brain proteomics comparing male and female *APOE* TR mice. A. Number of differentially expressed (DE) proteins for male vs. female mice. B. Volcano plot showing up and down regulated proteins in *APOE3* male vs. female mice. C. Volcano plot showing up and down regulated proteins in *APOE4* male vs. female mice. D. IPA of up and downregulated pathways in *APOE3* male vs. female mice. E. IPA of up and downregulated pathways in *APOE4* male vs. female mice. F. Venn diagram of upregulated protein overlap in *APOE3* and *APOE4* male vs. female mice. G. Venn diagram of down regulated protein overlap in *APOE3* and *APOE4* male vs. female mice.

**Supplemental Figure 2.**
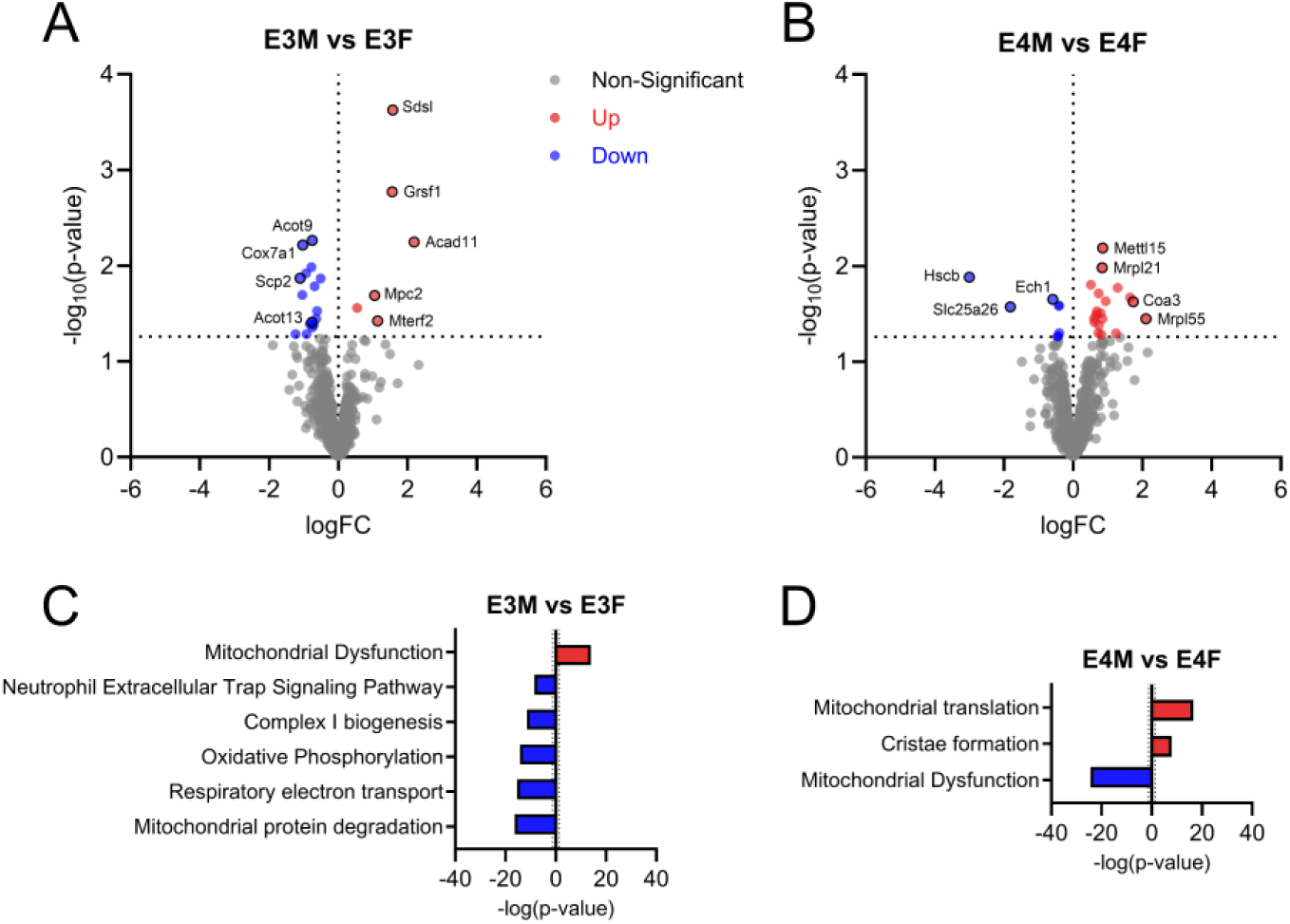
Isolated brain mitochondria proteomics comparing male and female APOE TR mice. A. Volcano plot showing up and down regulated mitochondrial proteins between *APOE3* male and female mice. B. Volcano plot showing up and down regulated proteins between *APOE4* male and female mice. C. IPA showing up and down regulated pathways between *APOE3* male and female mice. D. IPA showing up and down regulated pathways between *APOE4* male and female mice.

**Supplemental Figure 3.**
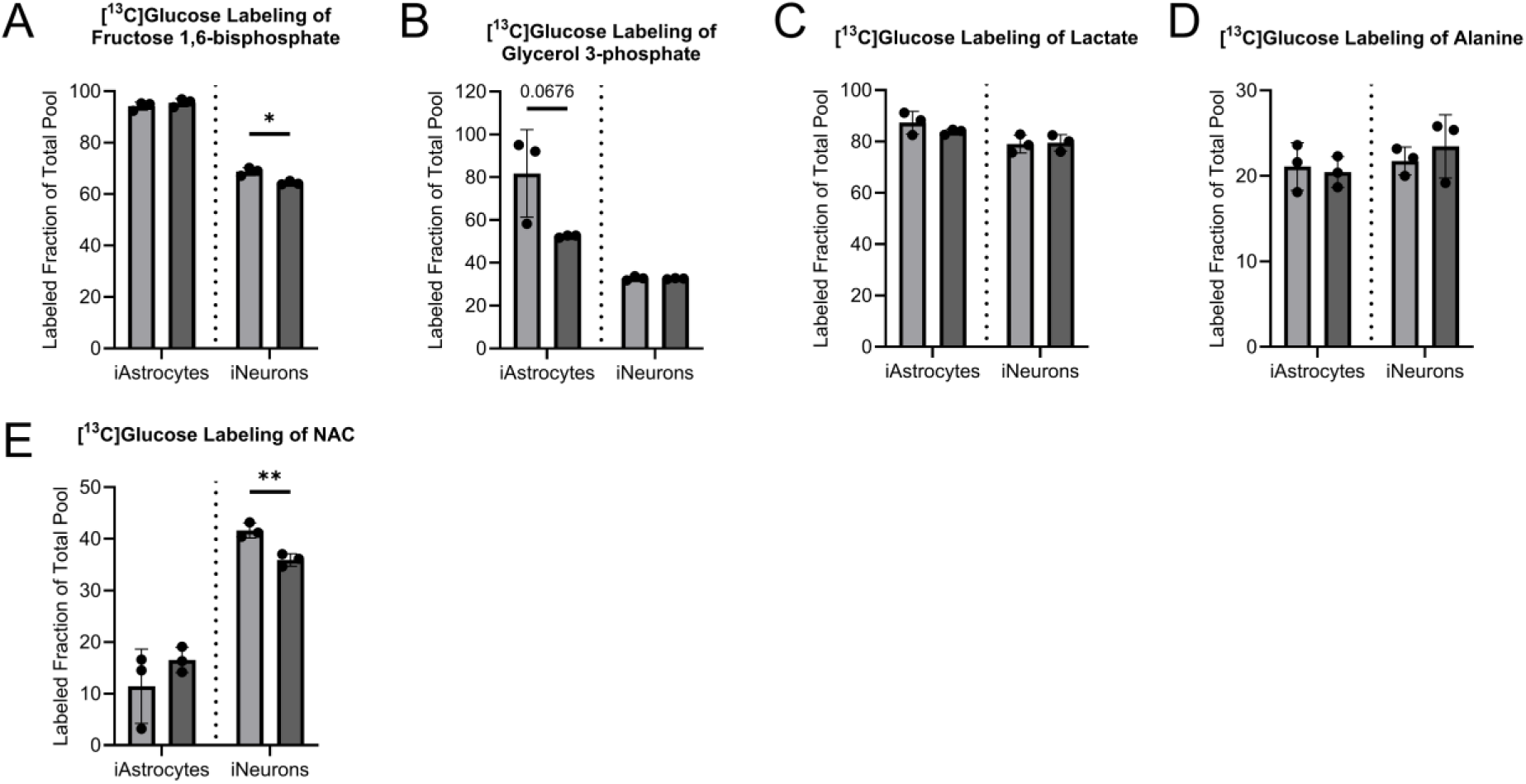
Stable isotope labeling of additional metabolites in iAstrocytes and iNeurons. iAstrocyte and iNeuron ^13^C-glucose labeling of A. fructose 1,6-bisphosphate, B. glycerol 3-phosphate, C. lactate, D. alanine, E. NAC. All data are shown as mean ±SD. *p<0.05, **p<0.01.

